# Genome-wide mutagenesis identifies factors involved in *Enterococcus faecalis* vaginal adherence and persistence

**DOI:** 10.1101/2020.04.30.069468

**Authors:** Norhan Alhajjar, Anushila Chatterjee, Brady L. Spencer, Lindsey R. Burcham, Julia L. E. Willett, Gary M. Dunny, Breck A. Duerkop, Kelly S. Doran

## Abstract

*Enterococcus faecalis* is a Gram-positive commensal bacterium native to the gastrointestinal tract and an opportunistic pathogen of increasing clinical concern. *E. faecalis* also colonizes the female reproductive tract and reports suggest vaginal colonization increases following antibiotic treatment or in patients with aerobic vaginitis. Currently, little is known about specific factors that promote *E. faecalis* vaginal colonization and subsequent infection. We modified an established mouse vaginal colonization model to explore *E. faecalis* vaginal carriage and demonstrate that both vancomycin resistant and sensitive strains colonize the murine vaginal tract. Following vaginal colonization, we observed *E. faecalis* in vaginal, cervical and uterine tissue. A mutant lacking endocarditis- and biofilm-associated pili (Ebp) exhibited a decreased ability to associate with human vaginal and cervical cells *in vitro*, but did not contribute to colonization *in vivo*. Thus, we screened a low-complexity transposon (Tn) mutant library to identify novel genes important for *E. faecalis* colonization and persistence in the vaginal tract. This screen revealed 383 mutants that were underrepresented during vaginal colonization at 1, 5 and 8 days post-inoculation compared to growth in culture medium. We confirmed that mutants deficient in ethanolamine catabolism or in the type VII secretion system were attenuated in persisting during vaginal colonization. These results reveal the complex nature of vaginal colonization and suggest that multiple factors contribute to *E. faecalis* persistence in the reproductive tract.

**IMPORTANCE:** Despite increasing prevalence and association of *E. faecalis* with aerobic vaginitis, essentially nothing is known about the bacterial factors that influence *E. faecalis* vaginal colonization. We have adapted an animal model of vaginal colonization that supports colonization of multiple *E. faecalis* strains. Additionally, we determined that ethanolamine utilization and type VII secretion system genes contribute to vaginal colonization and persistence. Identification of factors important for vaginal colonization and persistence provides potential targets for the development of therapeutics. This study is the first to identify key determinants that promote vaginal colonization by *E. faecalis*, which may represent an important reservoir for antibiotic resistant enterococci.

## INTRODUCTION

*Enterococcus faecalis* is an opportunistic pathogen that resides in the human gastrointestinal and urogenital tracts (1, 2). While *E. faecalis* colonization is normally asymptomatic, certain populations are at risk for severe disease including urinary tract infections (3), wound infections, pelvic inflammatory disease (PID), infective endocarditis, and adverse birth effects during pregnancy (reviewed in 4, 5). Enterococcal infections are often associated with the production of biofilms, assemblages of microbes enclosed in an extracellular polymeric matrix that exhibit cell-to-cell interactions (reviewed in 6). These biofilms have been observed on catheters, diabetic ulcers, and wounds resulting in severe infection. Treatment of enterococcal infections is becoming increasingly problematic due to their augmented ability to acquire mobile genetic elements, resulting in increased resistance to antibiotics, including “last-line-of-defense” antibiotics such as vancomycin (reviewed in 7, 8). Recently, there has been an increase in the emergence of vancomycin resistant enterococci (9), putting immunocompromised individuals at risk for developing severe chronic enterococcal infections. The emergence of vancomycin resistant enterococci (10) and its prevalence in both community and nosocomial settings is concerning and necessitates the development of alternative therapeutics to treat enterococcal infections.

*E. faecalis* encodes a multitude of virulence factors that allow the bacterium to colonize and persist in different sites of the human body. Surface proteins such as the adhesin to collagen (Ace), enterococcal fibronectin binding protein A (EfbA), aggregation substance (AS), and the endocarditis-and biofilm-associated pilin (Ebp) have been previously shown to play important roles in infective endocarditis and UTIs (reviewed in 11). Secreted factors such as gelatinase (12), autolysin A (13), and serine protease (SprE) are biofilm-associated factors that are involved in the degradation of host substrates, including collagen, fibrin and certain complement components (14). Many of these virulence factors are regulated via quorum sensing, which may be responsible for the switch from a commensal to pathogenic state (15-17).

Certain risk factors are associated with the transition of *E. faecalis* from commensalism to pathogenicity such as immune status, prolonged hospital stay, and the use of antibiotics (18). *E. faecalis* colonization and infection is often polymicrobial and these interactions have been observed in the intestine, bloodstream, and wounds (reviewed in 19). Furthermore, *E. faecalis* is frequently found in the vaginal tract of healthy women (20, 21) and its prevalence is increased in women diagnosed with aerobic vaginitis (AV), an inflammatory response accompanied by depletion of commensal *Lactobacillus sp*. and increased presence of opportunistic pathogens such as *E. faecalis*, Group B *Streptococcus* (GBS), *Staphylococcus aureus*, and *Escherichia coli* (22, 23). Symptoms of AV include malodor and discomfort, but AV can transition to more serious complications such as PID, severe UTIs, and complications during pregnancy. While it is evident that *E. faecalis* colonizes the human vaginal tract, the molecular determinants that allow enterococci to colonize and persist in the vaginal tract remain to be identified.

In this study, we modified our previously established GBS vaginal colonization model to analyze *E. faecalis* vaginal colonization and persistence. We determined that *E. faecalis* OG1RF (a rifampicin and fusidic acid derivative of strain OG1) and vancomycin resistant *E. faecalis* V583 can colonize and persist in the vaginal tract of CD1 and C57BL/6 mouse strains. We detected fluorescent *E. faecalis* in the vaginal lumen as well as the cervical and uterine tissues of colonized mice. Further, we demonstrated that an *E. faecalis* strain lacking Ebp pili is less adherent to vaginal cervical epithelium *in vitro*, but not attenuated *in vivo*. Thus, we screened an *E. faecalis* OG1RF transposon (Tn) mutant library for mutants that are underrepresented in the vaginal tract compared to the culture input, revealing multiple factors for *E. faecalis* persistence within the vagina. These factors include sortase-dependent proteins (SDPs), ethanolamine utilization genes, and genes involved in type VII secretion system (T7SS) machinery. We confirmed that a mutant strain in ethanolamine catabolism was significantly attenuated in the ability to colonize the vaginal epithelium, and T7SS was required to ascend in the female reproductive tract. This work is an important first step in identifying factors required for enterococcal vaginal colonization and will provide insight into potential therapeutic targets aimed at mitigating *E. faecalis* vaginal colonization in at-risk individuals.

## RESULTS

### *E. faecalis* colonization of the female reproductive tract

To characterize the ability of *E. faecalis* to interact with the epithelial cells of the lower female reproductive tract, we performed *in vitro* quantitative adherence assays using *E. faecalis* strains V583 (24) and OG1RF (25). An inoculum of 10^5^ CFU/well (multiplicity of infection [MOI] = 1) was added to a confluent monolayer of immortalized human vaginal and endocervical epithelial cells. Following 30 minutes of incubation, the cells were washed to remove non-adherent bacteria, the epithelial cells were detached from the plates, and adherent bacteria were plated on agar. Both strains exhibited substantial adherence to both cell lines (Fig. 1A). Next, we assessed the ability of *E. faecalis* to establish colonization of the murine vaginal tract. The vaginal lumen of C57BL/6 were swabbed and swabs were plated on CHROM™ agar to determine the presence of native enterococci. While native enterococci are detected on CHROM™ agar, no mice were colonized with strains that resemble *E. faecalis* V583 or OG1RF, as no colonies appeared on agar supplemented with antibiotics that select for V583 and OG1RF. Next, C57BL/6 mice were treated with β-estradiol 1 day prior to inoculation with 10^7^ CFU of *E. faecalis* V583 or OG1RF. After 1 day post-inoculation, the vaginal lumen was swabbed and bacteria were plated to enumerate *E. faecalis* V583 and OG1RF vaginal colonization levels (Fig. 1B). Swabs were plated on selective agar to ensure quantification of only the enterococcal strains of interest, restricting growth of native enterococcus. To determine whether *E. faecalis* ascends into reproductive tissues during colonization, murine vaginal, cervical, and uterine tissues were collected and homogenized to enumerate *E. faecalis* V583 abundance. *E. faecalis* was recovered from all mice 1 day post-inoculation in all tissues tested (Fig. 1C) and the CFU recovered from the vaginal swabs were similar to the total CFU counts from the vaginal tissue homogenates (Fig. 1B, C). This level and range of recovered *E. faecalis* CFU is similar to what we have observed using this mouse model for GBS and *S. aureus* vaginal colonization (26, 27). To visualize *E. faecalis* within the murine reproductive tract, mice were inoculated with either WT *E. faecalis* V583 or a V583 strain expressing green fluorescent protein (GFP) (28). These strains both colonize the vaginal tract (Fig. S1A). We harvested the female reproductive tract 1 day post-inoculation to avoid the loss of the GFP plasmid and made serial sections of these tissues, and performed fluorescent microscopy to visualize *E. faecalis*. We observed numerous fluorescent bacteria in the vaginal and uterine lumen (Fig. 1D, F) and embedded in the cervical lamina propria (Fig. 1E). We did not observe background green fluorescence in naïve mice (Fig. S1B, C, D), which coincides with previous experiments performed with GBS and *S. aureus* (26, 27). The presence of fluorescent *E. faecalis* in the cervix and uterus shows that *E. faecalis* can move from the vaginal lumen to the superior organs of the female reproductive tract.

**Figure 1:**
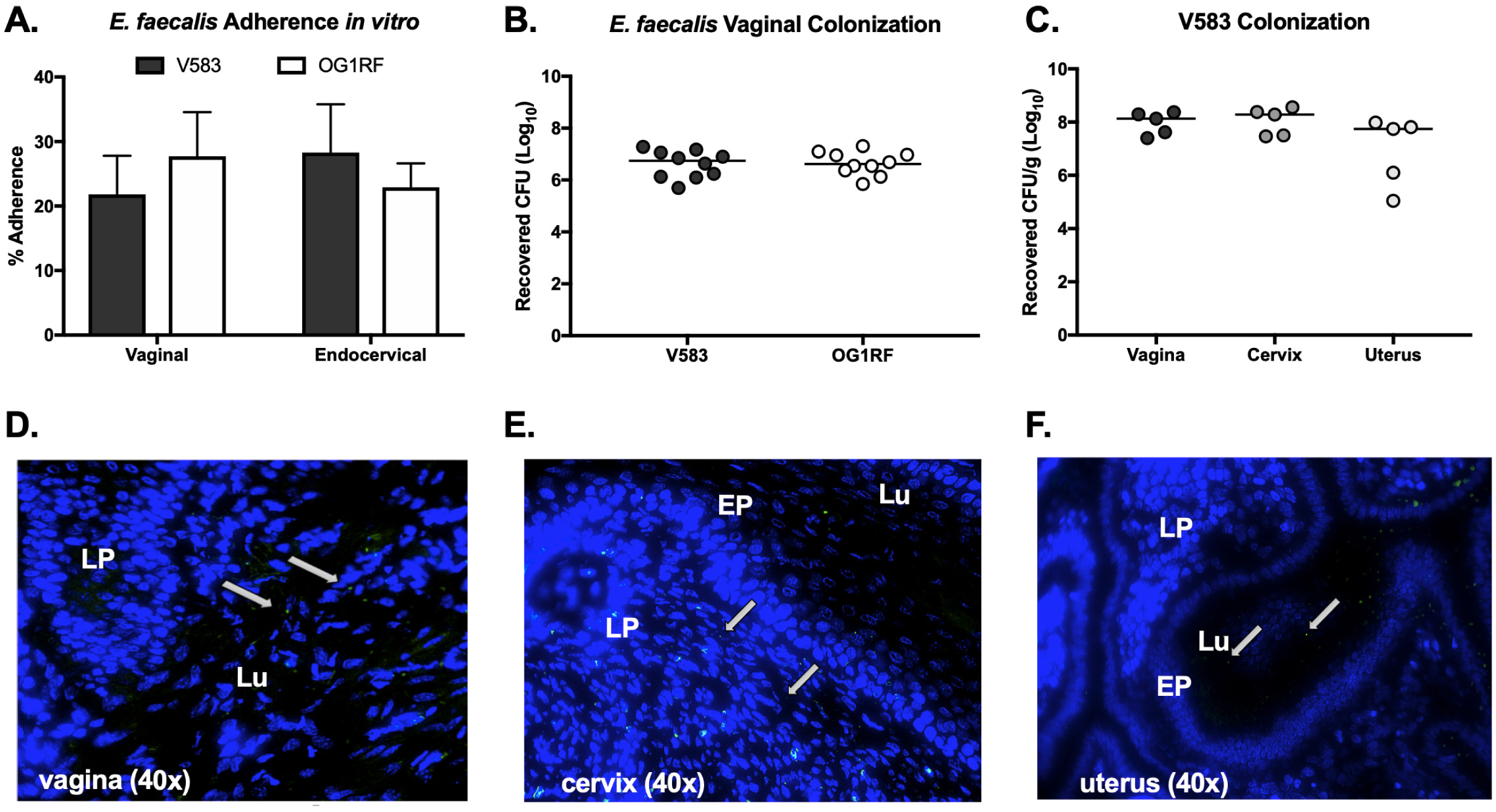
*E. faecalis* colonizes the murine female reproductive tract. (A) Adherence of *E. faecalis* V583 and OG1RF to human vaginal and endocervical cells. Data are expressed as percent recovered cell-associated *E. faecalis* relative to the initial inoculum. Experiments were performed in triplicate and error bars represent standard deviations (SDs); the results of a representative experiment are shown. (B) CFU counts of V583 and OG1RF recovered from vaginal swabs 1 day post-inoculation. (C) CFU counts of V583 from vaginal, cervical and uterine tissue 1 day post-inoculation. (D, E, F) Mice were inoculated with V583 expressing *gfp* and 7μm sections of vaginal (D), cervical (E) and uterine (F) tissue were collected 1 day post-inoculation and stained with DAPI for fluorescence microscopy. White arrows indicate green fluorescent bacteria present in tissue sections. Images were all taken at 40x magnification. LP = lamina propria, EP = epithelial layer, Lu = lumen.

### *E. faecalis* persists in the vaginal tract

To assess vaginal persistence, C57BL/6 mice were colonized with *E. faecalis* V583 or OG1RF and swabbed to determine bacterial load over time. Mice were swabbed daily until no colonies appeared on agar selective for V583 or OG1RF, indicating bacterial clearance from the vaginal lumen. While V583 persisted longer in the mouse vaginal tract, the mean CFU recovered for both V583 and OG1RF remained constant for the first week and then declined as mice eventually cleared both strains by 11-13 days (Fig. 2A, B). To determine if enterococcal vaginal persistence differs across mouse strains, C57BL/6 and CD1 mice were inoculated with V583 and swabbed over time. Both mouse strains were initially colonized with V583, but bacteria in C57BL/6 mice persisted longer (Fig. 2C, D). By day six only 20% of CD1 mice remained colonized compared to 85% of C57BL/6 mice. Differences in vaginal persistence may be due to differences in the native vaginal microbiota or immune status between mouse strains. It is also possible that bacteria occupy different niches within the reproductive tract of different mouse strains, which warrants further investigation. Overall, these results show that mouse strain background influences *E. faecalis* vaginal colonization and that C57BL/6 mice are a sufficient model to assess prolonged *E. faecalis* vaginal colonization and persistence.

**Figure 2:**
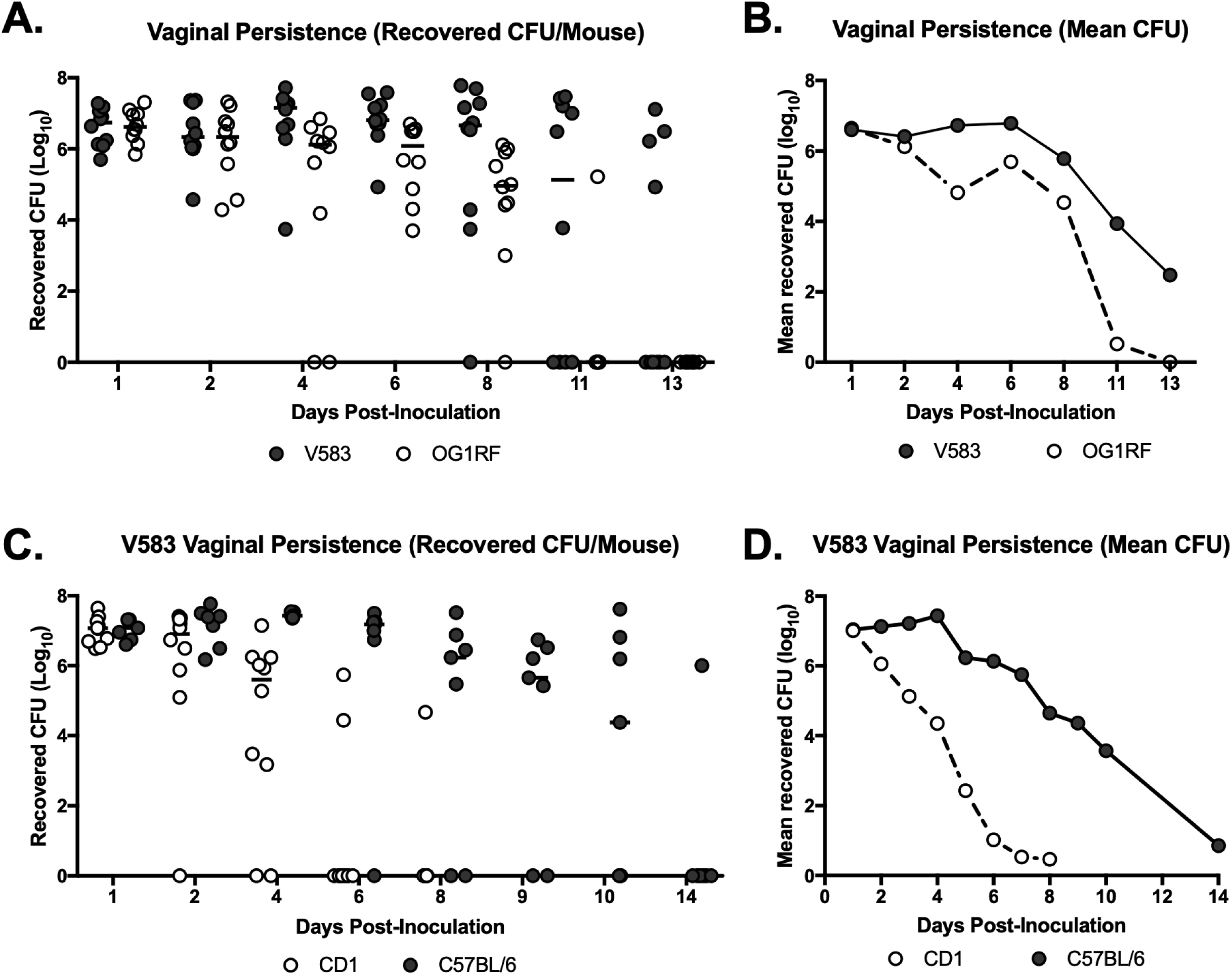
*E. faecalis* persists in the murine vaginal tract. (A and B) *E. faecalis* V583 and OG1RF in the murine vaginal tract. C57BL/6 mice (n = 10) were inoculated with 10^7^ V583 or OG1RF CFU and the vaginal lumen of each mouse was swabbed daily and swabs were serially diluted and plated on selective media to quantify CFU. Data are presented as recovered CFU per mouse (A) and mean recovered CFU (B). Data was analyzed using a Two-way ANOVA; ∗∗ P *<* 0.001. (C and D) CD1 (n = 10) and C57BL/6 (n = 7) mice were inoculated with V583 and the vaginal lumen of each mouse was swabbed daily, serially diluted, and plated on selective media to quantify CFU. Data are presented as recovered CFU per mouse (C) and mean recovered CFU (D). Black lines indicate the median of CFU values.

### Enterococcal pili contribute to interaction with reproductive tract tissues

The <u>e</u>ndocarditis- and <u>b</u>iofilm-associated <u>p</u>ilin (Ebp) of *E. faecalis* mediates infective endocarditis and UTIs(29-32), thus we hypothesized that Ebp may similarly contribute to vaginal colonization. To determine whether Ebp is important for facilitating interaction with the vaginal epithelium, we used a deletion mutant of *E. faecalis* OG1RF lacking all pilin structural components (Δ*ebpABC*) (33). We observed that the pilus mutant exhibited significantly reduced adherence to human vaginal and endocervical cells *in vitro* (Fig. 3A, B). To determine if Ebp is important for *in vivo* vaginal colonization and persistence, mice were inoculated with either WT OG1RF or OG1RF Δ*ebpABC* and colonization was quantified over the course of 12 days. We observed no differences in the CFU recovered from the vaginal lumen between WT OG1RF and OG1RF Δ*ebpABC* strains (Fig. 3C). Taken together these data suggest that Ebp contributes to *E. faecalis* attachment to reproductive tract tissues, but additional factors are likely required for persistence in the vaginal lumen *in vivo*.

**Figure 3:**
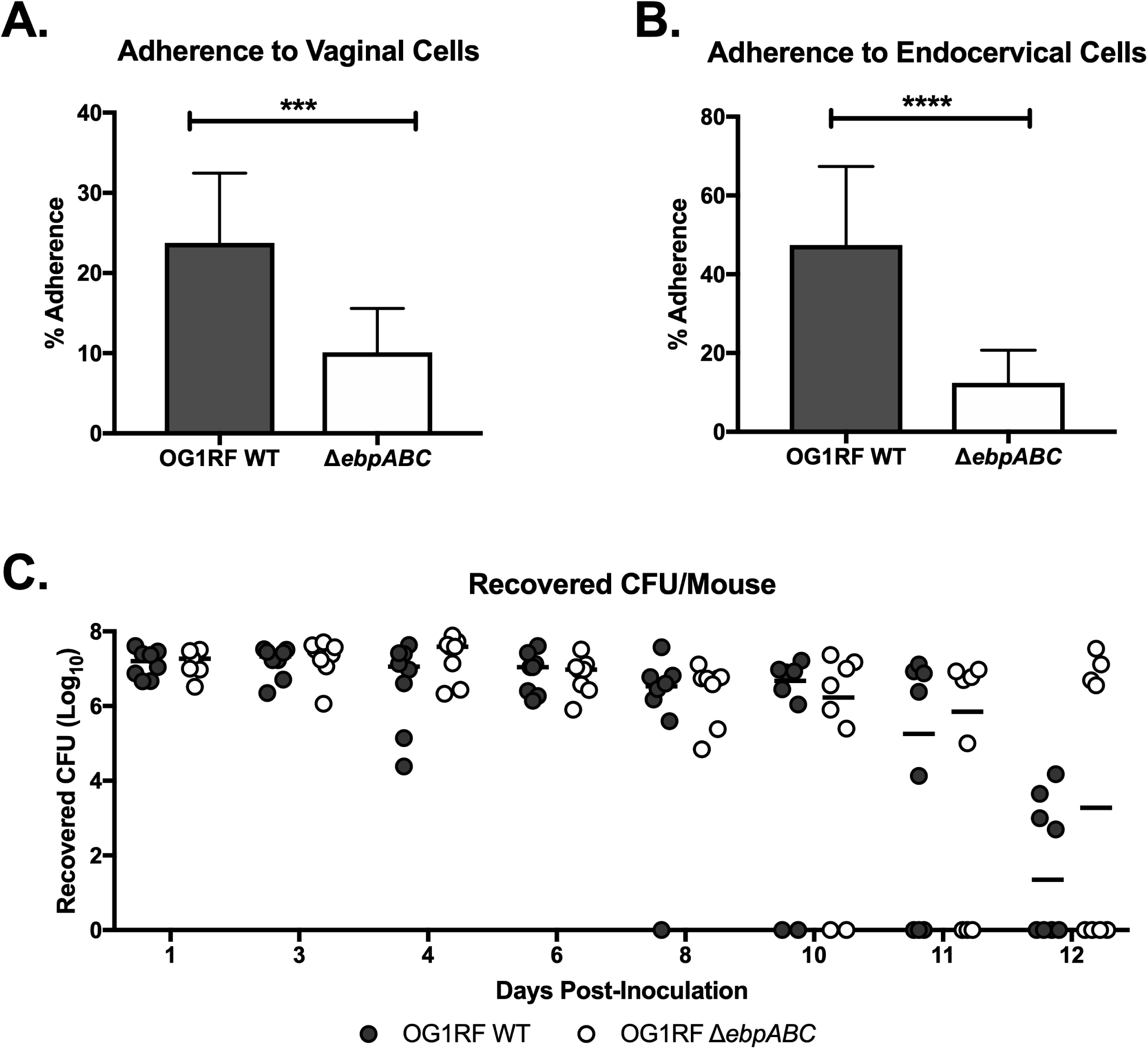
The role of enterococcal pili during vaginal colonization. (A and B) *E. faecalis* OG1RF WT and OG1RF Δ*ebpABC* adherence to human vaginal (A) and endocervical (B) cells. Data are expressed as percent recovered cell-associated *E. faecalis* relative to the initial inoculum. Experiments were performed with four technical replicates and error bars represent SDs; the results of three combined biological replicates are shown and analyzed using an unpaired t-test; ∗∗∗ P *<* 0.0001; ∗∗∗∗ P *<* 0.00001. (C) C57BL/6 mice were inoculated with either OG1RF WT or OG1RF Δ*ebpABC* and the vaginal lumen was swabbed daily. Data was analyzed using a Two-way ANOVA with Sidak’s multiple comparisons (P>0.05). Black lines indicate the median of CFU values.

### Identification of additional vaginal colonization factors by transposon mutagenesis analysis

To identify genetic determinants that confer enterococccal vaginal persistence, we used <u>s</u>equence-defined *<u>mar</u>iner* technology <u>tr</u>ansposon <u>seq</u>uencing (SMarT TnSeq) to screen an *E. faecalis* OG1RF Tn library consisting of 6,829 unique mutants (34). The library was grown to mid-log phase in triplicate and 10^7^ CFU of each replicate was vaginally inoculated into a group of 5 C57BL/6 mice (Fig. 4A). Vaginal swabs were plated on selective media daily for 8 days and CFU were quantified to assess colonization of the OG1RF Tn library compared to WT OG1RF (Fig. 4A and B). Genomic DNA was isolated from pooled Tn libraries recovered on days 1, 5, and 8 post-inoculation and Tn insertion junctions in *E. faecalis* genomic DNA were sequenced as described by Dale *et al*. (34). Sequenced reads were mapped to the *E. faecalis* OG1RF genome to identify genes that are necessary for *E. faecalis* vaginal colonization.

**Figure 4:**
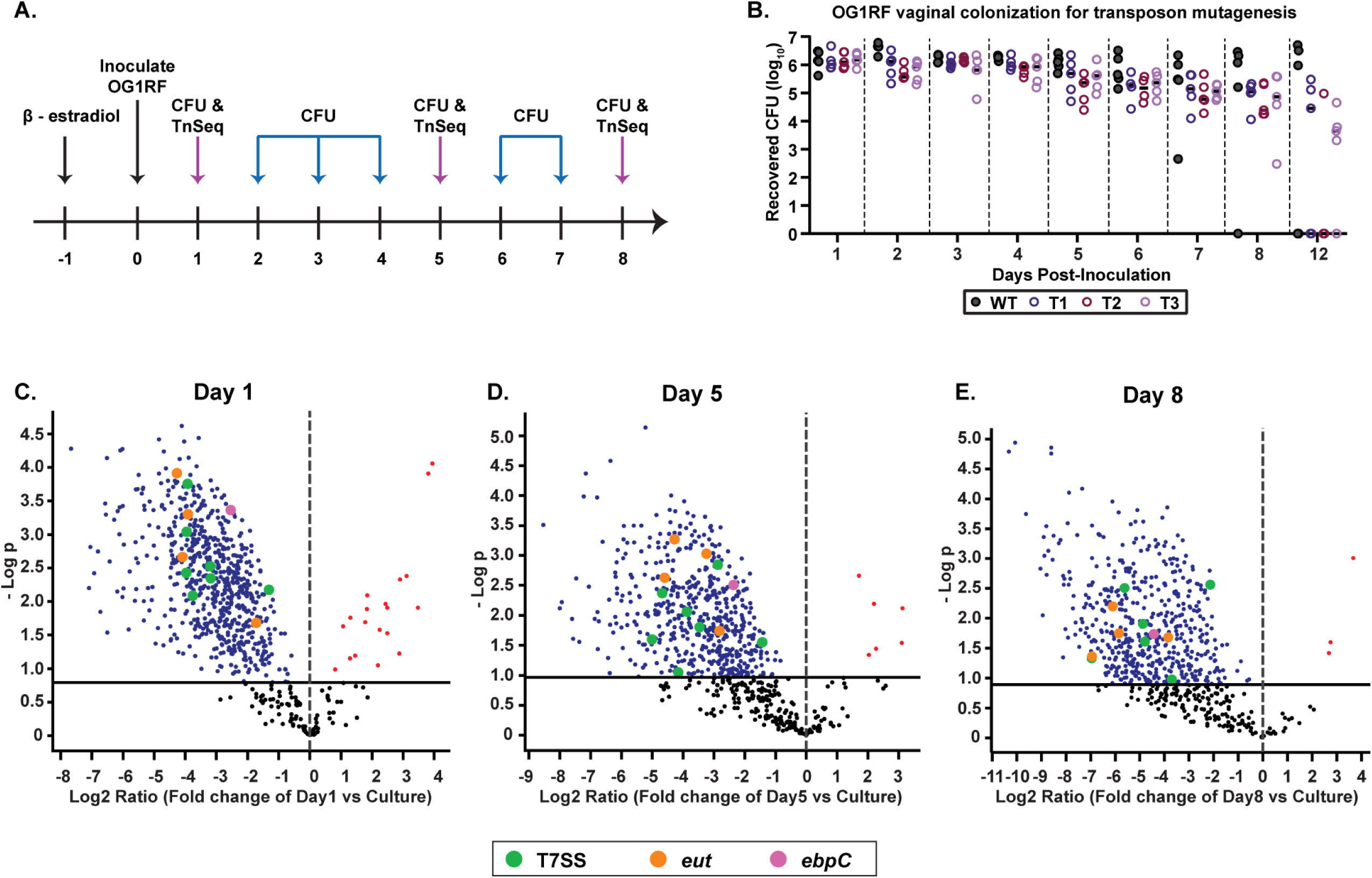
Identification of additional factors required for vaginal colonization and persistence by transposon mutant library screen. (A) Schematic representing experimental approach. C57BL/6 mice were treated with 17β-estradiol prior to inoculation with the OG1RF Tn library in triplicate groups or WT OG1RF. The vaginal lumen was swabbed and CFU was enumerated daily. DNA from recovered OG1RF Tn mutants was sequenced on days 1, 5 and 8. (B) CFU recovered from vaginal swabs of triplicate groups of mice colonized with OG1RF Tn mutagenesis library (T1, T2, T3) and OG1RF WT (n=5 mice per group). (C, D and E) Volcano plots depicting underrepresented and overrepresented mutants *in vivo* compared to culture on day 1 (C), day 5 (D) and day 8 (E) post-inoculation. (C) 667 underrepresented genes and 21 overrepresented genes *in vivo* compared to culture. (D) 404 underrepresented genes and 6 overrepresented genes *in vivo* compared to culture. (E) 507 underrepresented genes and 3 overrepresented genes *in vivo* compared to culture. Colored dots represent mutants of interest. Pink = *ebpC* Tn mutant, Orange = *eut* Tn mutants, Green = T7SS gene Tn mutants. Black solid line represents cut-off for statistical significance.

We observed that the *in vivo* vaginal environment altered the abundance of select mutants from the Tn library pool compared to the original culture input (Fig. 4C, D, E) (Tables S1 - S5). At day 1, a total of 667 depleted mutants were identified (Table S1), along with 544 (Table S2) and 507 (Table S3) at days 5 and 8 respectively; 383 of these mutants were identified at all three time points (Fig. 5A, Table S4A). Classification by clusters of orthologous groups of proteins (COGs) could be identified in 196 mutants from all 3 time points, the majority of which are involved in carbohydrate, amino acid, lipid and nucleotide transport/metabolism, as well as those involved in transcription and defense mechanisms (Fig. 5B). Of the remaining 187 mutants, 85 had insertions in intergenic regions and 102 were not assigned a COG domain. Interestingly the *ebpC* transposon mutant (*OG1RF_10871*, which encodes the shaft component of Ebp) was underrepresented at all time points (Fig. 4C, D, E, Table S4A). Additional mutants of interest included those involved in ethanolamine catabolism and T7SS components, in which various components of these systems were underrepresented at all three time points (Tables 1 and S4A). Furthermore, mutants for multiple sortase-dependent proteins (SDPs), including *ebpC*, were underrepresented at all time points (Tables 1 and S4A), suggesting that these factors may play important roles vaginal colonization and persistence.

**Table 1:** Selected list of differentially abundant transposon mutants during vaginal colonization compared to *in vitro* cultures.

| old_locus_tag | NCBI_description | Difference<br>(D1 - Cul) | Difference<br>(D5 - Cul) | Difference<br>(D8 - Cul) |
| --- | --- | --- | --- | --- |
| **Sortase-dependent proteins (SDPs)** |  |  |  |  |
| OG1RF_10811 | collagen adhesion protein | -4.33 | -4.14 | -5.72 |
| OG1RF_10871 | cell wall surface anchor family protein, *ebpC* | -2.54 | -2.37 | -4.44 |
| OG1RF_11531 | glycosyl hydrolase | -3.04 | -3.15 | -4.25 |
| OG1RF_11764 | cell wall surface anchor family protein | -2.80 | -3.48 | -5.24 |
| OG1RF_12054 | cell wall surface anchor family protein | -2.62 | -1.50 | -6.01 |
| **Ethanolamine utilization** |  |  |  |  |
| OG1RF_11342 | ethanolamine utilization protein EutL | -3.92 | -3.25 | -6.10 |
| OG1RF_11343 | ethanolamine ammonia-lyase small subunit | -4.09 | -4.60 | -5.85 |
| OG1RF_11344 | ethanolamine ammonia-lyase large subunit | -4.27 | -4.28 | -6.94 |
| **Type VII secretion system** |  |  |  |  |
| OG1RF_11103 | YukD superfamily protein, *esaB* | -3.19 | -5.02 | -6.95 |
| OG1RF_11109 | putative LXG-containing toxin | -3.93 | -3.47 | -4.87 |
| OG1RF_11111 | putative T7SS toxin | -3.76 | -4.15 | -4.80 |
| OG1RF_11113 | putative T7SS toxin | -3.19 | -2.87 | -3.70 |
| OG1RF_11114 | putative immunity protein | -3.96 | -3.88 | -5.62 |
| OG1RF_11122 | immunity protein | -1.31 | -1.43 | -2.14 |

**Figure 5:**
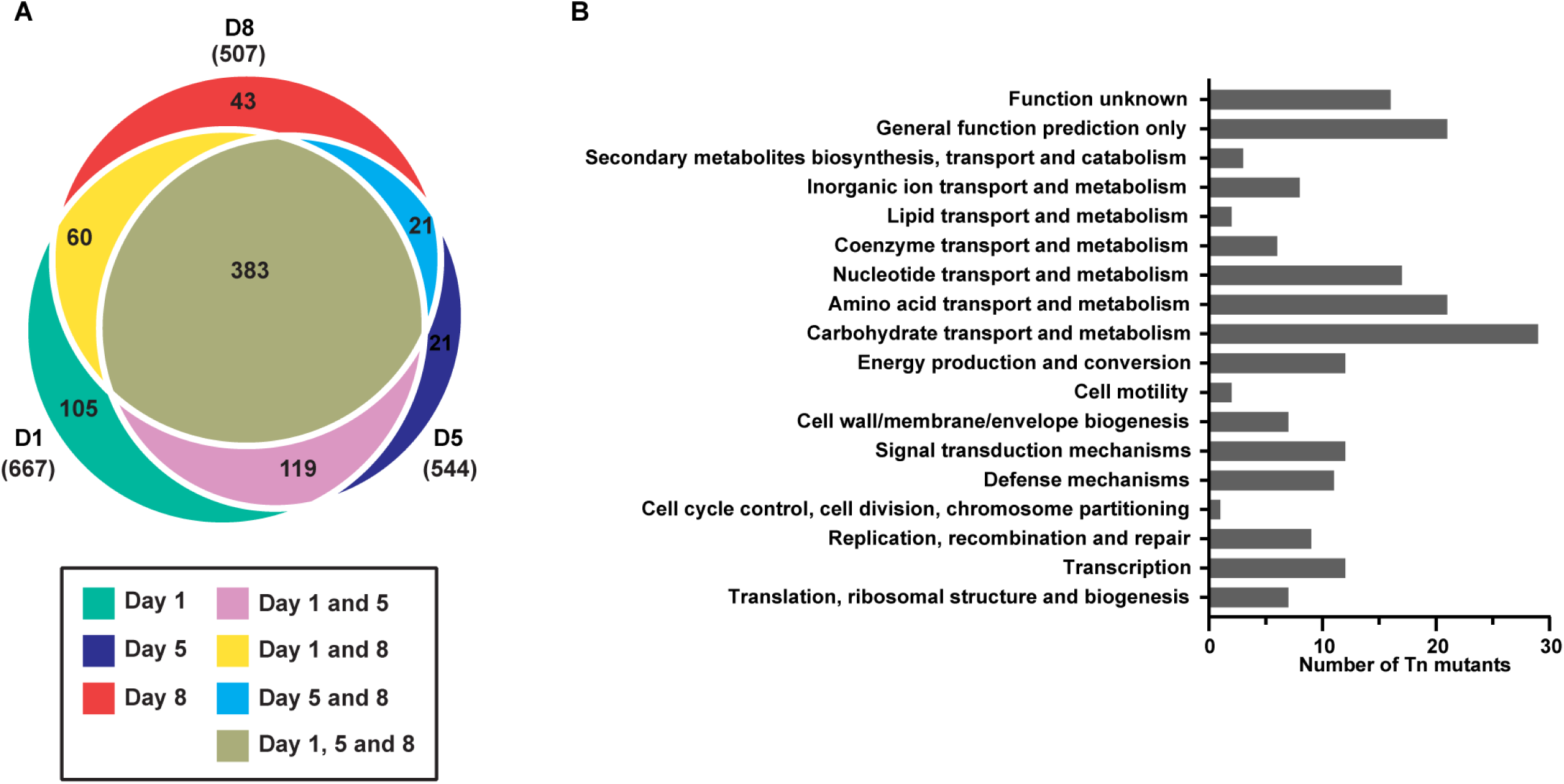
Classification of transposon insertion mutants by Cluster of Orthologous Groups (COGs). (A) Euler plot representing number of underrepresented mutants at all time points. (B) Cluster of Orthologous Groups (COGs) from underrepresented mutants common to all three time points categorized into functional categories. Total = 196 represents all common mutants that were assigned a COG domain.

Potential gain-of-function mutations have also been discovered during genome-wide library screens of fitness determinants in other bacteria (35-39). In addition to mutations that adversely impact vaginal colonization, our data shows that Tn insertions in 11 protein coding genes and 11 intergenic regions potentially enhance bacterial fitness *in vivo*. Nineteen of the 22 enriched mutants were common between days 1 and 5 whereas the other 3 were unique to day 8 (Table S5A and S5B). Since the majority of the mutants with increased fitness encode hypothetical proteins, the relationship between these genes and vaginal colonization is currently unclear and requires further investigation.

### Ethanolamine utilization and T7SS genes contribute to *E. faecalis* persistence in the reproductive tract

Our TnSeq analysis revealed many potential mutants that exhibited decreased colonization in the murine vaginal tract. We sought to confirm these results by analyzing mutants from systems with multiple affected genes. One significantly affected operon was ethanolamine (EA) utilization (*eut*) which consists of 19 genes in *E. faecalis* (40). Mutants in 4 *eut* genes were significantly underrepresented *in vivo* compared to the culture input at all time points. These included transposon mutants of the genes encoding both subunits for ethanolamine ammonia lyase, *eutB (OG1RF_11344)* and *eutC (OG1RF_11343)* (41), a carboxysome structural protein, *eutL (OG1RF_11342)*(42), and the response regulator, *eutV (OG1RF_11347)*, of the two-component system involved in the regulation of EA utilization (43) (Table S4A). To assess the importance of EA utilization on *E. faecalis* vaginal colonization, we co-colonized mice with *E. fae*calis OG1SSp (a derivative of OG1 that is resistant to streptomycin and spectinomycin) and an OG1RF Δ*eutBC* mutant (44). We note that both WT strains, OG1RF and OG1SSp, were able to colonize the vaginal tract at similar levels (Fig. S2). Further chromosomal DNA sequence comparison of OG1RF and OG1SSp revealed multiple nucleotide polymorphisms (SNPs) in OG1SSp, but no SNPs in genes in the *eut* locus (Table S6). Compared to the WT OG1SSp strain, the Δ*eutBC* mutant was cleared significantly faster from the mouse vagina as seen by CFU from individual mouse swabs, the mean CFU recovered and competitive index (CI) over time (Fig. 6A, B, C). These results suggest that the utilization of ethanolamine is important for enterococcal persistence in the vaginal tract.

**Figure 6:**
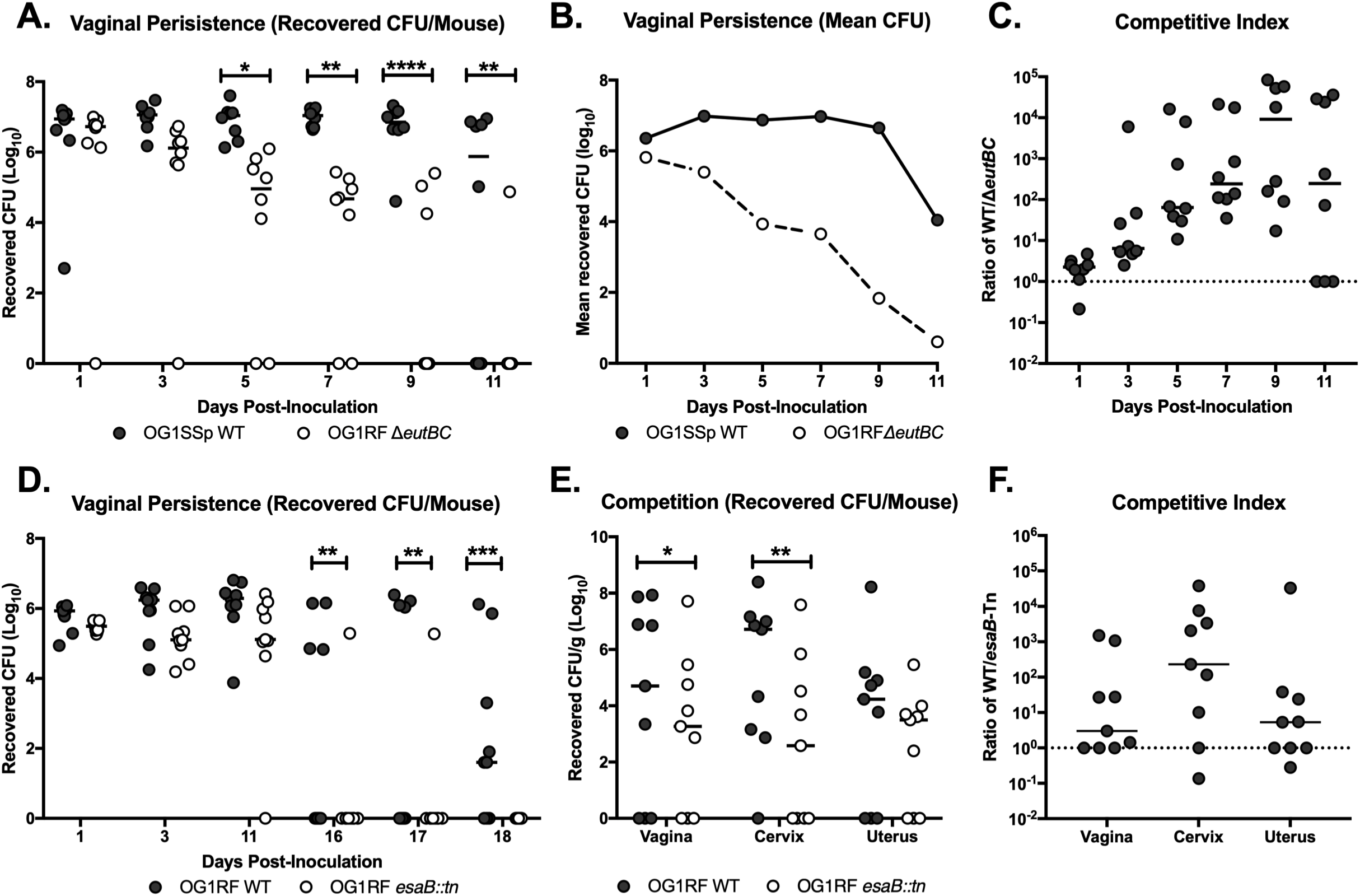
Ethanolamine utilization and type VII secretion system genes contribute to enterococcal persistence in the vaginal tract. (A, B and C) C57BL/6 mice were co-inoculated with OG1SSp WT and OG1RF Δ*eutBC* and vaginal lumen was swabbed to quantify CFU. Data are presented as recovered CFU per swab (A), mean recovered CFU (B) and CI between WT and mutant strains (C). Data was analyzed using a Two-way ANOVA with Sidak’s multiple comparisons; *∗* P < 0.05, *∗∗* P < 0.005, *∗∗\*\** P < 0.00005. (D) C57BL/6 mice were co-inoculated with OG1RF WT and OG1RF *esaB::tn* and vaginal lumen was swabbed to quantify CFU. Data was analyzed using a Two-way ANOVA with Sidak’s multiple comparisons; *∗∗* P < 0.005, *∗*∗* P < 0.0005 (E and F) C57BL/6 mice were co-inoculated with OG1RF WT and OG1RF *esaB*::*tn* and reproductive tissue was collected at 11 days post-inoculation. Data are presented as recovered Log_10_ CFU/gram (E) and CI between WT and Tn-mutant strain (F). CI is enumerated by calculating the ratio of WT to mutant *E. faecalis* recovered from the mouse reproductive tract. A CI >1 indicates an advantage to WT *E. faecalis*. Values below the limit of detection were enumerated as one-half the limit of detection. Data was analyzed using a paired t-test; *∗* P < 0.05, *∗∗* P < 0.005. Black lines indicate the median of CFU values.

We also observed that Tn insertion mutants in the T7SS locus were significantly underrepresented at all time points *in vivo* compared to the culture input. The T7SS has been shown to play an important role in virulence in multiple bacterial species such as *Staphylococcus, Listeria* and *Bacillus* (45). In *E. faecalis*, genes in the T7SS locus have been shown to be induced during phage infection (46). We observed that transposon mutants for *esaB* (*OG1RF_11103)*, a putative cytoplasmic accessory protein; *OG1RF_11109* and *OG1RF_11111*, putative toxin effector proteins; *OG1RF_11113*, a putative toxin, *OG1RF_11114*, a transmembrane protein; and *OG1RF_11122*, a potential antitoxin protein were all significantly underrepresented (Table S4A). To confirm the role of the T7SS in vaginal colonization, we utilized an *esaB* (*esaB::tn*) mutant, in which *esaB* is disrupted by a transposon and thus Tn-mediated polar effects may exist for this strain. Following co-colonization with *E. faecalis* OG1RF and the OG1RF *esaB::tn* mutant strain, we observed that while there were no differences in initial colonization, fewer *esaB::tn* mutant bacteria were recovered from the vaginal lumen at later time points (Fig. 6D). Since we only observed differences in colonization between WT OG1RF and OG1RF *esaB::tn* at later time points, we performed subsequent experiments to determine whether there were differences in ascension between the two strains. Mice were co-colonized with the two strains and we harvested tissues at day 11, before there were any colonization differences observed between the two strains. Here, we observed that WT OG1RF outcompeted the *esaB::tn* mutant strain and was better able to access reproductive tract tissues (Fig. 6E, F), indicating that the T7SS may be involved in vaginal persistence and ascension in the female reproductive tract.

## DISCUSSION

*E. faecalis* is associated with a wide spectrum of infections, particularly under immunocompromised states and during compositional shifts in the host microbiota (47, 48). Although an increasing body of evidence links enterococci with bacterial vaginosis (BV) and aerobic vaginitis (AV) (22, 23, 49-51), the molecular determinants that facilitate *E. faecalis* colonization and persistence in the vaginal tract are largely unknown. Here, we employed *in vitro* and *in vivo* systems to acquire genome-scale interactions that confer *E. faecalis* fitness within the female reproductive tract. We show that both vancomycin sensitive enterococci (VSE) and VRE adhere to cell types of vaginal and cervical origin, a signature of bacterial colonization that precedes tissue invasion and systemic spread. Further, genetic features involved in biofilm formation, ethanolamine utilization and polymicrobial interactions influence *E. faecalis* vaginal carriage.

Previous studies have demonstrated the importance of Ebp pili in enterococcal virulence and biofilm formation (19). We found that deletion of *ebpABC* attenuated binding to human vaginal and endocervical cells but did not influence bacterial burden in the vaginal lumen, similar to the observed function of Ebp pili in the intestine (52). Enhanced *E. faecalis* adherence in tissue culture compared to *in vivo* colonization may reflect the lack of liquid and mucus flow that bacteria encounter within the vaginal tract, emphasizing the significance of our animal model for investigating *E. faecalis*-vaginal interactions. Consistent with this observation, an *in vivo* Tn library screen revealed only two underrepresented biofilm-associated mutants, *ebpC-Tn* and *OG1RF_10506-Tn*, encoding a putative polysaccharide deacetylase homolog implicated in low biofilm formation *in vitro* (53, 54). Together, these results show that individual mutations in *ebp* or other well characterized biofilm genes are not sufficient to impair vaginal niche establishment and/or persistence of enterococci, which likely depends on the concerted effort of multiple factors. Furthermore, similar to *ebpC-Tn*, we observed that genes for multiple sortase-dependent proteins (SDPs) were underrepresented at all time points during vaginal colonization. The genome of OG1RF contains 21 sortase-dependent proteins, including Ebp (52). Other than *ebpC* (*OG1RF_10871*), we observed that 4 other SDPs are underrepresented during vaginal colonization, including *OG1RF_10811, OG1RF_11531, OG1RF_11764* and *OG1RF_12054* (Table 1). Previous reports indicate the importance of SDPs during gastrointestinal colonization by enterococci (52), implicating the possibility that multiple SDPs also play a role during vaginal colonization.

Transition from nutrient rich laboratory media to the vaginal tract likely imparts dramatic alterations in *E. faecalis* metabolism. In support of this hypothesis, our high-throughput Tn mutant screen showed that mutations in carbohydrate, amino acid and nucleotide metabolic pathways were indispensable in the vaginal tract. Specifically, we showed that WT bacteria outcompete a *eut* locus mutant during vaginal colonization. In contrast, Kaval and colleagues demonstrated that mutations in *eut* genes leads to a slight increase in fitness within the intestine (55). This observation likely reflects varying metabolic requirements of enterococci in different host environments. While a number of reports exist on the contributions of EA catabolism in host-bacteria interactions within the intestine (56), studies are lacking for the relevance of this EA metabolism in other host-associated environments. Our results raise important questions regarding EA utilization in the female reproductive tract. Although commensal microbes and the epithelium are rich sources of EA, the composition and source of EA in the vaginal tract remains unknown. A recent report showed that *E. faecalis* EA utilization attenuates intestinal colitis in IL10 knockout mice in the presence of a defined microbiota (57). Whether EA utilization promotes virulence or commensalism for enterococci in the context of vaginal tissue remains to be determined. Considering that the by-product of EA metabolism, acetate, is anti-inflammatory and promotes IgA production in the intestine (58, 59), it is intriguing to consider that enterococcal EA catabolism might modulate immune responses within the female reproductive tract.

T7SSs have been implicated in the maintenance of bacterial membrane integrity, virulence, and inter-bacterial antagonism (45, 60-66). In *S. aureus*, T7SS encoded proteins confer protection from membrane damage caused by host fatty acids (65, 66). Although *E. faecalis* T7SS genes were shown to be induced in response to phage driven membrane damage, direct contributions of these genes in cell envelop barrier function and/or virulence in the context of animal models are poorly defined. Our TnSeq analysis revealed that insertional mutations in six T7SS genes diminished early and late vaginal colonization by *E. faecalis* OG1RF. In vaginal co-colonization competition experiments, an *esaB::tn* strain reached WT colonization levels early and showed a defect in long term persistence. The incongruence in the colonization kinetics of T7SS mutant strains compared to T7SS-Tn library mutants which were observed early after inoculation, presumably stem from the inherent differences in the vaginal milieu in these two experiments. The TnSeq library employed in this screen is a complex population of approximately 7,000 unique mutants, and it is very likely that direct or indirect interactions between mutants influences fitness. LXG–domain toxins, which are part of the T7SS, have been shown to antagonize neighboring non-kin bacteria (63). The fact that two mutants with LXG–domain encoding gene mutations, *OG1RF_11109* and *OG1RF_11111*, were underrepresented across all time points during vaginal colonization suggests that these putative antibacterial proteins may influence enterococcal interactions with the resident microbes of the vaginal tract.

In addition to genes encoding SDPs, ethanolamine utilization and T7SS, other Tn mutants that were underrepresented in all time points are worth discussing. For example, *OG1RF_12241*, a homolog of the oxidative stress regulator *hypR*, was underrepresented at all time points (Table S4A). We have recently shown that this gene is involved in phage VPE25 infection of *E. faecalis* OG1RF (46). Furthermore, enterococcal mutants in the CRISPR/*cas9* locus (*OG1RF_10404* and *OG1RF_10407*) were underrepresented at all three time points (Table S4A). While the role of CRISPR-Cas systems in providing prokaryotic immunity to mobile genetic elements has been extensively investigated, there is also evidence suggesting that this system may be involved in other prokaryotic processes besides adaptive immunity. Cas9 has been shown to have various functions in regulation of virulence in a number of bacteria including *Francisella novicida, Campylobacter jejuni*, and *Streptococcus agalactiae* (67-69). In *E. faecalis*, CRISPR-Cas-harboring strains are associated with increased capacity to form biofilms and increased mortality in a mouse urinary tract model (70). Our Tn-seq analysis further reveals the potential importance for Cas9 during vaginal colonization, which warrants follow up studies.

While a majority of underrepresented mutants were common to all time points, we identified certain mutants unique to either early or late colonization. For example, the ethanolamine utilization protein EutQ (*OG1RF_11333*), a classified acetate kinase in *Salmonella enterica* (71), was significantly underrepresented at day 1, but not the later time points. We also observed that Tn mutants for the transmembrane signaling protein kinase IreK (*OG1RF_12384*) were underrepresented only at day 1. In *E. faecalis*, IreK is involved in regulation of cell wall homeostasis (72), long-term persistence in the gut and has also been shown to be essential for enterococcal T7SS expression and subsequent activity (73). These proteins may therefore be important contributors to enterococcal vaginal colonization, though further investigation is required. Our Tn-seq analysis also identified Tn mutants that were unique to day 8 post-inoculation, indicating that these factors may be important for later stage colonization and persistence. One observed mutant was the *fsrB* gene (*OG1RF_11528*) of the Fsr quorum-sensing system, which directly regulates virulence factors such as serine protease and gelatinase, while also indirectly regulating other virulence factors involved in surface adhesion and biofilm development (74-76). Although it is not well understood whether biofilms are being formed during vaginal colonization, certain hits in our Tn-seq (i.e. *ebpC* and *fsrB*) analysis suggests that biofilm-associated factors play a role in enterococcal persistence in the vaginal tract. Bacterial mutants for the response regulator *croR* (*OG1RF_*12535) were also underrepresented only at the later time point. CroR has shown to be involved in virulence regulation, cell wall homeostasis and stress response, and antibiotic resistance (77-79). Finally, underrepresentation of the sortase-associated gene (*OG1RF_10872*) was also unique to late colonization. The underrepresentation of enterococcal mutants late in vaginal colonization suggests these factors may be important for long-term persistence of *E. faecalis* in the vaginal tract. While the majority of underrepresented mutants were common to all time points, mutants unique to certain time points indicate that some factors may be important for either initial colonization or enterococcal survival in the vaginal tract.

Here, we report the utilization of a mouse model for investigating host-enterococcal interactions in the vaginal tract. This will be a useful model for analyzing the bacterial and host factors that govern enterococcal vaginal colonization, as well as characterizing the polymicrobial interactions that may contribute to *E. faecalis* niche establishment and persistence. Transposon library screening of *E. faecalis* recovered from the mouse vagina has revealed new insights into our understanding of enterococcal vaginal carriage. Our results emphasize the importance of ethanolamine utilization and T7SS components for successful *E. faecalis* colonization of the female reproductive tract, highlighting the complex nature of this niche.

## METHODS

### Bacterial strains and culture conditions

A detailed list of bacterial strains can be found in Table S7. *E. faecalis* strains V583 (80) and OG1-(RF and SSp) (70, 81) were used for these experiments. *E. faecalis* was grown in brain heart infusion (BHI (82)) broth at 37 °C with aeration and growth was monitored by measuring the optical density at 600nm (OD_600_). For selection of *E. faecalis* V583, BHI agar was supplemented with gentamicin (100 μg/ml). For selection of *E. faecalis* OG1RF, OG1RF Δ*ebpABC* (33) and OG1RF Δ*eutBC* (44), BHI agar was supplemented with rifampicin (50 μg/ml) and fusidic acid (25 μg/ml). For selection of *E. faecalis* OG1SSp, BHI agar was supplemented with streptomycin (150 μg/ml) and spectinomycin (100 μg/ml). *E. faecalis* OG1RF *esaB*::*tn* (34) was grown on BHI agar supplemented with rifampicin (50 μg/ml), fusidic acid (25 μg/L), and chloramphenicol (15 μg/ml).

### *In vitro* adherence assays

Immortalized VK2 human vaginal epithelial cells and End1 human endocervical epithelial cells were obtained from the American Type Culture Collection (VK2.E6E7, ATCC CRL-2616 and End1/E6E7, ATCC CRL-2615) and were maintained in keratinocyte serum-free medium (KSFM; Gibco) with 0.1 ng/mL human recombinant epidermal growth factor (EGF; Gibco) and 0.05 mg/ml bovine pituitary extract (Gibco) at 37°C with 5% CO^2^. Assays to determine cell surface-adherent *E. faecalis* were performed as described previously when quantifying GBS adherence (83). Briefly, bacteria were grown to mid-log phase (OD_600_ = 0.4 - 0.6) and added to cell monolayers (multiplicity of infection [MOI] = 1). After a 30 minute incubation, cells were washed with phosphate-buffered saline (PBS) three times following detachment with 0.1 mL of 0.25% trypsin-EDTA solution and lysed with addition of 0.4 mL of 0.025% TritonX-100 in PBS by vigorous pipetting. The lysates were then serially diluted and plated on Todd Hewitt agar (THA) to enumerate the bacterial CFU. Experiments were performed at least three times with each condition in triplicate, and results from a representative experiment are shown.

### Murine vaginal colonization model

Animal experiments were approved by the Institutional Animal Care and Use Committee at University of Colorado-Anschutz Medical Campus protocols #00316 and #00253 and performed using accepted veterinary standards. A mouse vaginal colonization model for GBS was adapted for our studies (26). Eight-week old female CD1 (Charles River) or C57BL/6 (Jackson) mice were injected intraperitoneally with 0.5 mg 17β-estradiol (Sigma) 1 day prior to colonization with *E. faecalis*. Mice were vaginally inoculated by gently pipetting 10^7^ CFU of *E. faecalis* in 10μL PBS into the vaginal tract, avoiding contact with the cervix. On subsequent days the vaginal lumen was swabbed with a sterile ultrafine swab (Puritan). For co-colonization, mice were inoculated with two of the following *E. faecalis* strains: OG1SSp, OG1RF or deletion mutants in the OG1RF background. To assess tissue CFU, mice were euthanized according to approved veterinary protocols and the female reproductive tract tissues were dissected and placed into 500μL PBS and bead beat for 2 min to homogenize the tissues. The resulting homogenate was serially diluted and *E. faecalis* CFU enumerated on BHI agar supplemented with antibiotics to select for the strain of interest.

### Histology

Mice were inoculated with *E. faecalis* V583 containing a plasmid that expresses *gfp* (pMV158GFP) and contains resistance to tetracycline (15 μg/mL) [23]. After 1 day post-inoculation, the murine female reproductive tract was harvested, embedded into Optimal Cutting Temperature (OCT) compound (Sakura), and sectioned at 7μm with a CM1950 freezing cryostat (Leica). For fluorescence microscopy, coverslips were mounted with VECTASHIELD mounting medium containing 4’,6-diamidino-2-phenylindole (DAPI, Vector Labs). Images were taken with a BZ-X710 microscope (Keyence) [22].

### Transposon mutant library growth and vaginal colonization

The *E. faecalis* OG1RF transposon mutant library was generated previously (34). The E. *faecalis* OG1RF pooled transposon library was inoculated into 5 ml of BHI at a total of 10^8^ CFU and grown with aeration to an OD_600_ of 0.5. The library was inoculated into the vaginal tracts of C57BL/6 mice at 10^7^ CFU. The library was plated on BHI agar to confirm the inoculum for all groups of mice. Mice were swabbed daily and swabs were plated on BHI supplemented with rifampicin (50 μg/ml), fusidic acid (25 μg/ml) and chloramphenicol (20 μg/ml) to quantify CFU. On days 1, 5, and 8, undiluted swabs were plated on BHI agar with antibiotics and grown to a bacterial lawn. Bacteria were scraped and re-suspended in PBS and pelleted. DNA from days 1, 5, 8, and the input culture was extracted from pellets using a ZymoBIOMICSTM DNA MiniPrep kit (Zymo Research).

### Transposon library sequencing and data analysis

Transposon-junction DNA library preparation and Illumina NovaSeq 6000 DNA sequencing (150 base paired end mode) was performed by the Microarray and Genomics Core at the University of Colorado Anschutz Medical Campus as previously described (46). For downstream analysis of transposon-junctions, we used only the R1 reads generated by paired end sequencing. Illumina adapter trimmed raw reads were mapped to the *E. faecalis* OG1RF reference sequence (NC_017316.1) and differentially abundant transposon mutants were identified using statistical analysis scripts established by Dale *et al*.(34). The abundance of Tn mutants in culture was compared to input library used for culture inoculation and mutants that are not significantly different (p > 0.05) between these two samples were considered for the next steps of the analysis. For comparisons between *in vivo* and *in vitro* samples, mutants were considered significantly different if the adjusted P value was < 0.05 and a log_2_ (fold change) > 1.

### Genomic DNA sequencing and comparative analysis

*E. faecalis* genomic DNA was purified using a ZymoBIOMICS(tm) DNA Miniprep Kit (Zymo Research) and 150 bp paired end sequencing was performed on Illumina NextSeq 550 by the Microbial Genome Sequencing Center, University of Pittsburgh. *E. faecalis* OG1SSp genome DNA was purified using a Qiagen DNeasy kit and was sequenced on the MiSeq platform (2 × 75 bp) at the University of Minnesota Genomics Center. All reads were mapped to *E. faecalis* OG1RF reference sequence (NC_017316.1) using CLC Workbench (Qiagen). The basic variant caller tool in CLC Genomics Workbench was used to identify single nucleotide polymorphisms using default settings (similarity fraction = 0.5 and length fraction = 0.8).

### Data availability

The Tn-Seq and genomic DNA reads have been deposited at the European Nucleotide Archive under accession numbers PRJEB37929 and PRJEB39171, respectively.

### Statistical analysis

GraphPad Prism version 7.0 was used for statistical analysis and statistical significance was accepted at P values of < 0.05 (∗ P < 0.05; ∗∗ P < 0.005; ∗∗∗ P < 0.0005; ∗∗∗∗ P < 0.00005). Specific tests are indicated in figure legends.

## ACKNOWLEGMENTS

We thank Kimberly Kline for providing *E. faecalis* OG1RF Δ*ebpABC*, Danielle Garsin for providing *E. faecalis* OG1RF Δ*eutBC*, Katrina Diener and Monica Ransom for custom TnSeq library preparation and the Microarray and Genomics Core at the University of Colorado Anschutz Medical Campus for DNA sequencing. This study was supported by the NIH 5T32AI007405-28 to B.L.S., the NIH/NIAID R21 AI130857 to K.S.D, the NIH/NIAID R01 AI141479 to B.A.D, and a University of Colorado School of Medicine IMPA to K.S.D and B.A.D.

