## Supplementary figures and images for "Genome-wide mutagenesis identifies factors involved in *Enterococcus faecalis* vaginal adherence and persistence"

### Supplemental Figure 1

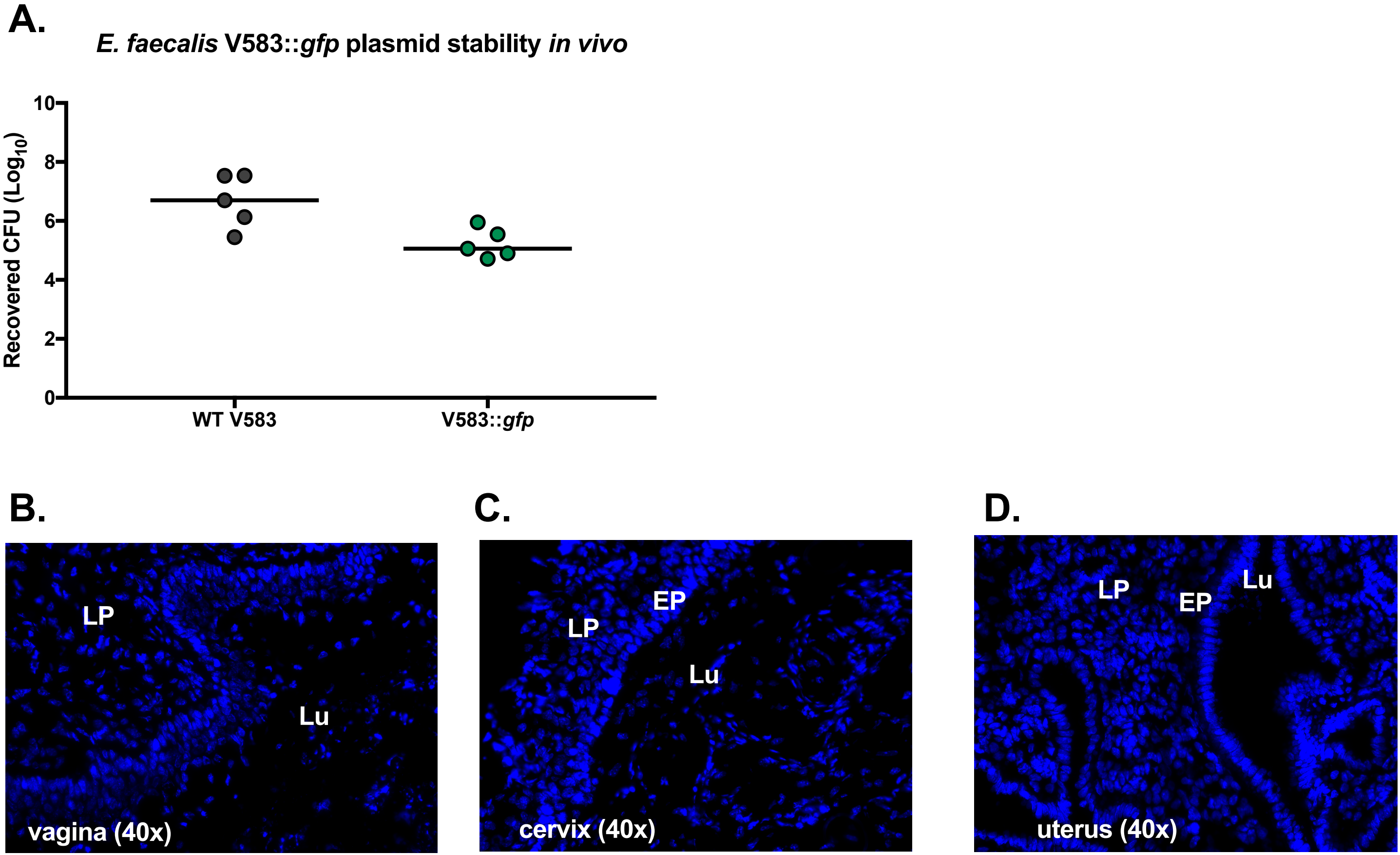

### Supplemental Figure 2

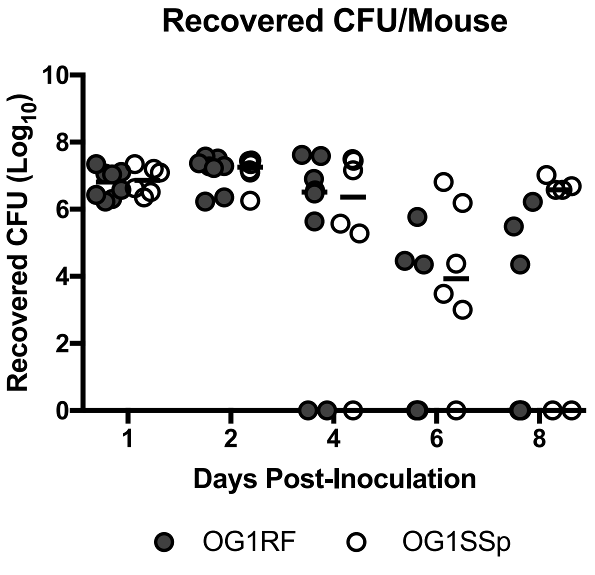
