## Supplemental Figure Legends for "Genome-wide mutagenesis identifies factors involved in *Enterococcus faecalis* vaginal adherence and persistence"

**Supplemental Figure 1: Colonization of *E. faecalis* V583::*gfp*.** (A) C57BL/6 mice were inoculated with either WT V583 or V583::pMV158GFP and the vaginal lumen was swabbed to quantify CFU. Data are presented as recovered CFU per swab. WT V583 was selected by plating on agar supplemented with gentamycin (100μg/ml) and V583::pMV158GFP was selected by plating on agar supplemented with gentamycin (100μg/ml) and tetracycline (15μg/ml) to determine GFP plasmid stability. (B, C, D) Naïve C57BL/6 mice were treated with β-estradiol and 7μm sections of vaginal (B), cervical (C) and uterine (D) tissue were stained with DAPI for fluorescence microscopy. Images were all taken at 40x magnification. LP = lamina propria, EP = epithelial layer, Lu = lumen.

**Supplemental Figure 2: Colonization and persistence of *E. faecalis* OG1- (RF and SSp).** C57BL/6 mice were co-inoculated with OG1RF WT and OG1SSp WT and the vaginal lumen was swabbed to quantify CFU. Data are presented as recovered CFU per swab. Data was analyzed using a Two-way ANOVA with multiple comparisons; $P > 0.05$.
